# Acute testicular hyperthermia leads to a rapid loss of global piRNA levels and a consequent increase in transcript abundance, including LINE1 activity within heat-sensitive male germ cells

**DOI:** 10.1101/2025.03.30.646213

**Authors:** Benjamin R. Robinson, Jacob K. Netherton, Rachel A. Ogle, Sean M. Burnard, Grace E. Williams, Georgia M. Tennant, Heather J. Lee, Mark A. Baker

## Abstract

**Objective:** To understand the immediate impact that testicular heat stress has on isolated populations of precursor male germ cells including Spermatocytes and Spermatids.

**Design:** Mice were given testicular heat stress and pre-cursor male germ cells were immediately isolated. RNA sequencing was performed and validated using qPCR.

**Subjects:** This work was carried out in adult male CD1 mice.

**Results:** Using next-generation RNA sequencing 134 differentially expressed transcripts were found to be differentially expressed upon exposure to testicular hyperthermia, 93% of which were upregulated. In addition, testicular hyperthermia induced 395 differential splicing events and altered the usage of 61 polyadenylation sites. To explain these observations and understand why transcript abundance appears to favour upregulation following testicular hyperthermia, we assessed global piRNA levels and found an overall, rapid reduction. Concomitantly, we observed an increase in transposable element RNA (LINE1) and protein (ORF1p) abundance. Furthermore, increased LINE1 expression appeared to be correlated with DNA damage in the male germline. At 24 hours post heat stress, piRNA levels recovered close to control levels, coincident with a significant reduction in LINE1 transcript expression within spermatocytes.

**Conclusion:** Testicular hyperthermia needs to be considered in the context of all reproductive outcomes. Affected spermatozoa are likely to be genetically compromised, leading to adverse outcomes such as infertility or loss of embryo following fertilization.

## Introduction

For healthy sperm to be produced spermatogenesis must occur 2-8°C below core body temperature[1–3]. Testicular hyperthermia, brought about through elevated air temperatures (35- 48.8°C)[4–7], scrotal insulation[8–15], cryptorchidism[16] or the use of a hot water bath[17] leads to the production of fewer sperm, a higher proportion of which are of poor quality[5–15, 17, 18]. In addition, spermatozoa produced during periods of heat stress present with increased levels of DNA damage. When these cells are used for fertilization, either through assisted conception or natural pregnancy, this contributes to higher rates of embryo loss[12, 19–22].

The adverse effects of testicular hyperthermia has its origins in precursor germ cells, the most sensitive of which are spermatocytes followed by round spermatids[23] (recently reviewed in [23]). As such, following testicular hyperthermia, sperm with abnormal morphology and impaired motility are observed in the ejaculate 14-30 days post intervention[24]. Importantly, prolonged or higher temperature exposures can lead to more severe outcomes such as complete azoospermia.

While the physiological and cellular responses to testicular hyperthermia are well documented, there is paucity of information when it comes to understanding the underlying biochemical mechanisms. To gain insight, several groups have opted for a transcriptomic approach using either microarrays[25–28] or next-generation RNA sequencing (NGS) [29–32]. Collectively, over 2700 transcripts have been reported to be differentially expressed, most of which are observed 24 hours post intervention[31]. However, gene expression changes have been documented to occur as early as 30 minutes post heat, and as late as 95 days[32]. In addition, specific alterations in non-coding RNA species including between 11-279 miRNAs[29–31], 172 circRNAs[31] and 465 lncRNAs[31] have been shown to be differentially expressed following testicular hyperthermia. While these studies provide valuable insight into the transcriptional effects of testicular heat stress, they do not fully explain the initial molecular mechanisms underlying testicular dysfunction. Additionally, many of these studies have used total testis RNA[30–32] which likely masks critical changes in heat sensitive cell types, such as round spermatids and spermatocytes.

Another class of non-coding RNA that has recently generated great interest are Piwi-interacting RNA (piRNA). These non-coding RNA species are enriched in the testis and play essential roles in spermatogenesis[33, 34], including the ability to supress Transposable Element (TE, formerly known as “Jumping Genes”) expression[35–37]. As retrotransposons, if TE activity is left unchecked, they have the capacity to insert into new genomic locations, causing widespread DNA damage, mutations and alter gene regulation[38–43]. Notably, 24 hours following a 15- minute hot water bath (43°C), 4,745 and 7,688 piRNA transcripts were up or downregulated respectivley in mouse testis[34]. Similarly, using a surgically cryptorchid rat model, 6.8% of piRNA transcripts were upregulated while 6.18% were downregulated three days post procedure[33]. This data suggests that piRNA are highly sensitive to hyperthermia, with some of the highest number of differentially expressed transcripts being documented to date. However, whether heat stress has an immediate impact on piRNA levels remains unknown.

Herein, we examine the early effects of testicular heat stress on mRNA, piRNA and TE expression initially within round spermatids, then compared specific responses to spermatocytes. Our data suggests that a global reduction in piRNA, which is more significant in spermatocytes than round spermatids, precedes the up regulation of many mRNA species, including TE. This increased activity of TE may explain the origins of sperm DNA damage and has particular implications for the assisted conception industry.

## Materials and Methods

### Key Resources Table

| REAGENT or Resource | SOURCE | IDENTIFYER |
| --- | --- | --- |
| **Antibodies** |  |  |
| Anti-ORF1p | Abcam | CAT# ab216324 |
| Anti-α-Tubulin | Sigma-Aldrich | CAT# T6074 |
| Rabbit anti-mouse HRP | ThermoFisher | CAT# A16166 |
| Goat anti-rabbit HRP | Sigma-Aldrich | CAT# DC03L |
| **Chemicals** |  |  |
| Hoechst 33342 | Sigma-Aldrich | CAT#B2261 |
| Propidium Iodide | Sigma-Aldrich | CAT#P4170-25MG |
| Collagenase type I | Sigma-Aldrich | CAT#C1-28-100MG |
| DNase I | Sigma-Aldrich | CAT#10104159001 |
| Trypsin-EDTA solution 0.25% | Sigma-Aldrich | CAT#T4049 |
| DMEM, high glucose | ThermoFisher | CAT#11965092 |
| RNeasy Plus Mini kit | Qiagen | CAT#74136 |
| TUBO DNA-free kit | ThermoFisher | CAT#AM1907 |
| QuantiTect Reverse Transcription kit | Qiagen | CAT#205311 |
| PowerUp SYBR Green Master Mix | ThermoFisher | CAT#A25742 |
| Click-iT Plus TUNEL assay-Alexa Fluor 647 | ThermoFisher | CAT#C10619 |
| **Deposited Data** |  |  |
| Round spermatid RNA seq Fasta files | This paper | Complete dataset is available at Gene Expression Omnibus accession no. GSE290301 |
| **Experimental models: Organisms/strains** |  |  |
| Mus Musculus Swiss (CD1) | Australian BioResources | Swiss CD1 |
| **Software and algorithms** |  |  |
| TEtranscripts | Hammell Lab, Cold Spring Harbour | Jin, Y., et al., (2015) [44] |
| DESeq2 R package | Bioconductor | RRID:SCR_015687 |
| Interactive Genomics Viewer | Broad Institute | RRID:SCR_011793 |
| MAGIQ | Github | Vaquero-Garcia, J., et al. (2016)[45] |
| QAPA | Github | Ha, K.C., B.J. Blencowe, and Q. Morris. (2018)[46] |
| ClusterProfiler R package | Bioconductor | RRID:SCR_016884 |
| GraphPad Prism | GraphPad | RRID:SCR_002798 |

### Animals

Adult male Swiss (CD1) mice aged 8-14 weeks were held in standard animal housing conditions with free access to food and water. All experimental procedures undertaken were approved by the University of Newcastle Animal Care and Ethics Committee under the approval number A2019-921 and A20.

### Testicular heat stress model

Testicular heat stress was induced by anesthetizing mice (5% isoflurane - 2L/minute) and placing them on a polystyrene floating block with a hole, allowing the scrotum to be submerged in a thermally controlled water bath, but little to no other part of the mouse. Anesthesia was maintained via nose cone (1.5-2% isoflurane - 2L/min). Water temperature was maintained at 42°C or 33°C for heat and control groups respectively. After 30 minutes of exposure, mice were removed from the water bath, dried, allowed to recover from anesthesia then immediately sacrificed by CO2 asphyxiation at the times indicated.

### Analysis of Epididymal Spermatozoa

The procedure for the removal of spermatozoa from the epididymis is shown here [47]. For morphology analysis, sperm were fixed in 2% paraformaldehyde (ThermoFisher) and then evaluated under phase contrast microscopy. At least 100 cells were assessed for each biological sample.

### Purification of round spermatids and pachytene spermatocytes

Following heat stress, testes were decapsulated and seminiferous tubules manually dissociated. Interstitial cells were removed by a two-step enzymatic digestion. Tubules were incubated for 20 min at 33°C in high glucose Dulbecco’s Modified Eagle Medium (ThermoFisher) containing 0.5mg/ml collagenase type I (Sigma-Aldrich), 0.5mg/ml DNase (Sigma-Aldrich), washed, then incubated for an additional 20 min at 33°C in high glucose DMEM containing 0.25% (w/v) Trypsin/EDTA (Sigma-Aldrich), 0.5mg/ml DNase. Digestion was halted by addition of 0.25% (w/v) Bovine Serum Albumin. Cells were washed, resuspended in DMEM, then passed through a 70µm nylon cell strainer to form a single cell suspension. Cell concentrations were determined by hemocytometer. For NGS, round spermatids were isolated by FACS as described by [48] and [49] with minor modifications. For all other isolation of round spermatids and spermatocytes, STA-PUT velocity sedimentation was used as described by [50]. Herein, the single cell suspension was loaded onto a 2-4% (w/v) BSA, high glucose DMEM gradient, and 5 ml fractions were collected after 3.5 hours of sedimentation at 25 °C. Fraction purity was calculated by counting cell types based on nuclear morphology (<100 cells/fraction) after Hoechst 33342 staining, fixation with 4% paraformaldehyde and visualizing with a Zeiss Axio observer fluorescent microscope.

### RNA isolation and RT-qPCR

RNA was extracted from isolated cell populations using the RNeasy Plus Mini kit (Qiagen) and residual genomic DNA was removed with the TUBO DNA-free kit (ThermoFisher). For RT-qPCR, 1µg RNA was reverse-transcribed into cDNA using the QuantiTect Reverse Transcription kit with random hexamer primers (Qiagen) according to manufacturer instructions. RT-qPCR was performed using PowerUp SYBR Green Master Mix (ThermoFisher) on a QuantStudio 5 Real-Time PCR system with standard cycling conditions, 60°C annealing temperature and 500nM primer concentration (N=≥3, samples ran in triplicate). Relative gene expression was quantified using the 2^-ΔΔCT^ method [51] with β-actin as reference gene. Internal control primers designed for regions of consensus across alternative isoforms were also used as reference genes for confirmation of LSV events. Primer sequences are provided in supplementary table S5 (LINE1 primer sequences are sourced from [52]).

### Whole transcriptome sequencing and differential expression analysis

Total RNA was extracted using RNeasy mini-RNA kit (Qiagen) and sent to Australian Genome Research facility (AGRF) for Illumina NovaSeq 150bp paired end total transcriptome RNA sequencing (N=5). For analysis of differential gene and TE expression, cleaned sequence reads were mapped to genomic sequences (*Mus musculus* GRCm38) using STAR aligner. TEtranscripts[44] was used to generate gene and TE counts. Lowly expressed genes and TEs, defined as ≥5 counts in ≥5samples, were excluded for further analysis. Differential expression was analysed using the DESeq2 R package[53], filtering for features with base-mean count >10, FDR >0.05, and absolute fold change >1.5. Gene ontology enrichment analysis was performed with the enrichGO function of the clusterProfiler R package[54].

### Local splice variation and alternative ploy(A) analysis

Cleaned sequence reads were mapped to reads were mapped to the Mus musculus mm39 genome using the STAR aligner by AGRF. The MAGIQ software package[45] was used to detect and quantify LSV events (Δψ>0.1, confidence interval >0.9). Sashimi plots of LSV events were generated using Interactive Genomics Viewer (IGV)[55]. Alternative polyadenylation analysis was performed using QAPA[46] (padj>0.05).

### Immunoblotting

Total protein was extracted from heat stressed and control spermatocytes (N=3) by lysis and homogenisation in 100µl SDS extraction buffer (10ml stock; 2ml 10% w/v sodium dodecyl sulfate (Sigma-Aldrich), 1g sucrose (Sigma-Aldrich), 1x cOmplete mini protease inhibitor tablet (Roche), 2.5ml 0.375M Tris pH 6.7, 938µl 1M Tris pH 7.5, 10mM TCEP (Sigma-Aldrich), up to 10ml distilled water). Protein lysates were separated by 4-20% SDS-PAGE gel (Bio-Rad) electrophoresis and transferred to a PVDF membrane. Membranes were bloked in TBST wash buffer (4.8g Tris base (Astral), 17.6 sodium chloride (Sigma-Aldrich), 1% (w/v) Tween-20 (Astral), pH 7.6)), containing 3% (w/v) BSA (Sigma-Aldrich), for 1 hour at 25°C, then incubated overnight at 4°C with primary antibody against ORF1p 1:1000 (Abcam ab216324) or α-Tubulin (Sigma-Aldrich T6074) in 3% BSA-TBST . Membranes were washed and incubated with HRP-conjugated 1:2500 anti-rabbit (ThermoFisher A16166) or 1:3000 anti-mouse (Sigma-Aldrich DC03L) in 3% BSA-TBST for 1 hour at 25°C. Chemiluminescent detection was performed with ECL prime detection reagent (Cytiva) and imaged using a ChemiDoc MP Imaging system (Bio-Rad). ORF1p band intensity was scaled and normalised to scaled α-tubulin band intensity using Fiji image analysis software.

### Denaturing Urea PolyAcrylamide Gel Electrophoresis (Urea-PAGE)

For piRNA resolution and quantification, RNA was extracted with TRIzol reagent (ThermoFisher) and was separated using 15% acrylamide urea-PAGE (4.8g Urea (Sigma Aldrich), 3.75ml 40% (19:1) acrylamide/Bis (Bio-Rad), 15µl 30% ammonium persulfate (Sigma-Aldrich), 2µl TEMED (Sigma-Aldrich), 5ml TBE buffer (10.6g Tris base (Astral), 5.5g Boric acid (Sigma Aldrich), 0.75g EDTA (Sigma Aldrich), up to 1L distilled water), distilled water up to 10mL).

Prior to loading, 500ng RNA sample was made up to 10µl in 50% formamide (v/v) (Sigma Aldrich), heated to 70°C for 10 min, then returned to ice prior to loading. Gels were run at 200V for 90min then stained with with GelRed nucleic acid stain (Millipore) for 15 min. Gels were imaged using a ChemiDoc MP Imaging system (Bio-Rad). Relative piRNA levels were calculated by normalising to the average scaled intensity of 6 control RNA bands using FIJI image analysis software.

### TUNEL assay

Testes were fixed in 4% paraformaldehyde (ThermoFisher) overnight at 4°C, then imbedded in paraffin for sectioning. Sections (5µm) were rehydrated through sequential xylene and graded ethanol washes. Heat-induced antigen retrieval was performed by boiling for 20 minute boil in sodium citrate buffer (2.9g sodium citrate (Sigma Aldrich), 500µl Tween-20 (Astral), up to 1L distilled water, pH 6). DNA damage was detected using the Click-iT Plus TUNEL assay-Alexa Fluor 647 (ThermoFisher) according to manufacturers instructions. A negative control was prepared by omitting the terminal deoxynucleotidyl transferase (TdT) enzyme. Sections were counterstained with 1:10000; 10mg/ml Hoechst 33342 (Sigma-Aldrich) to stain nuclei, washed, then mounted with Mowiol antifade medium. Images were captured using a Zeiss Axio observer fluorescent microscope with an Axiocam 305 mono camera and processed using Fiji Image analysis software. The percentage of TUNEL-positive cells was calculated as the number of TUNEL positive cells/ total cell number of Hoechst 33342 positive cells x 100 (N=3).

### Statistical significance

Data was analysed and graphed using GraphPad Prism 10.1 with data represented as mean ± SEM. Data normality was determined using the Shaprio-Wilk test and statistical significance assessed by unpaired student’s t tests.

## Results

### Increased production of poor-quality spermatozoa following testicular hyperthermia

To assess the impact of a single testicular heat event on spermatogenesis and validate our model, we exposed the testis of mice to a 42°C water bath for 30 minutes. Consistent with others, 3 weeks post heat we observed a significant decrease in total and progressive (Fig. 1a/b) motility, together with an increase in sperm displaying morphological defects (Fig. 1c/d). With respect to the latter, testicular heat stress led to the increased production of sperm with poor head shape (Fig. 1d, bottom left), head and tail defects (Fig. 1d, top right) and poor acrosome formation (Fig. 1d, bottom right) compared to control (Fig. 1d, top left).

**Figure 1.**
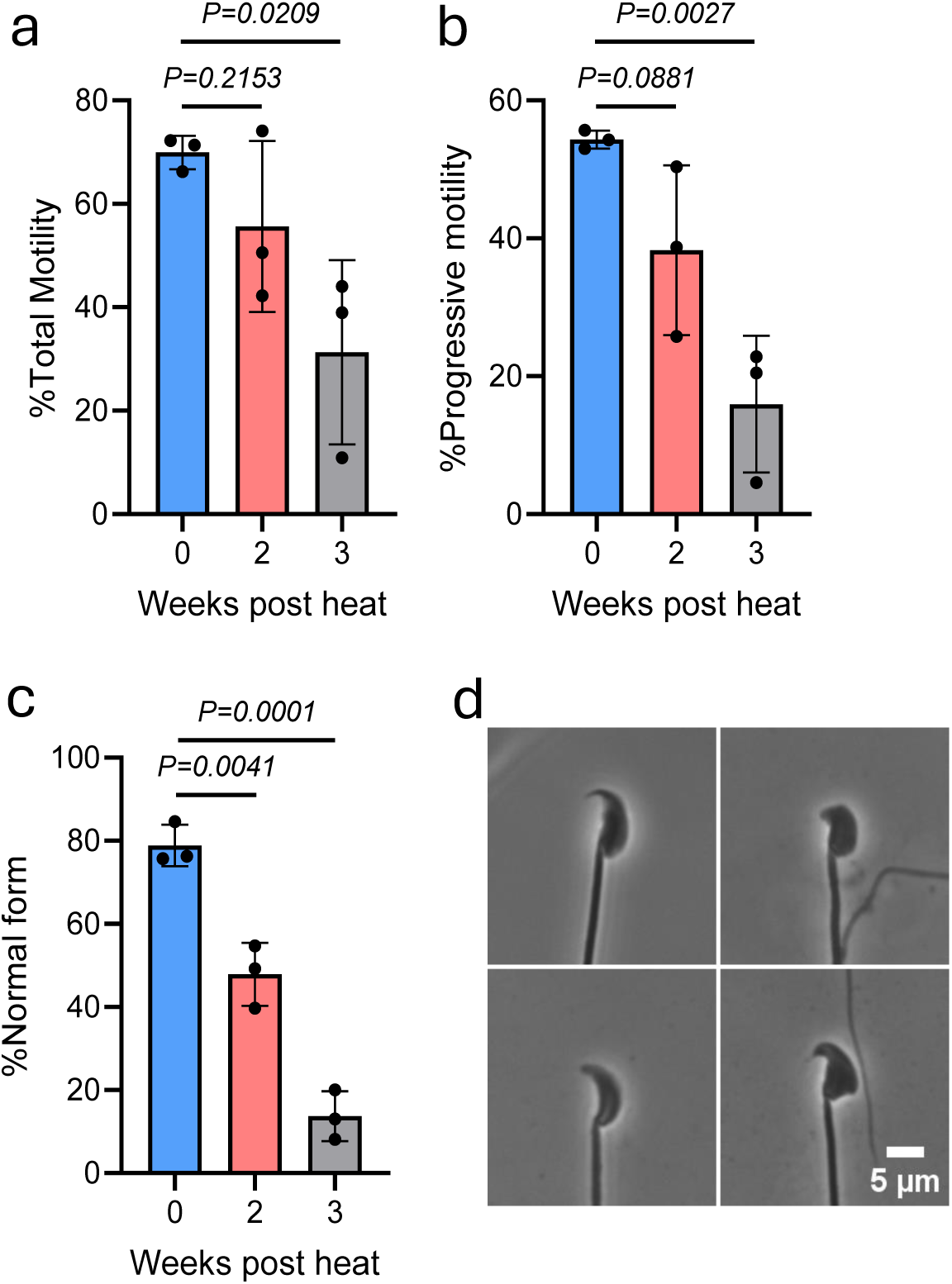
Increased production of poor-quality spermatozoa following testicular hyperthermia. Mice were given either testicular heat stress (42°, 30 mins) or anaesthetic control (33°C, 30 mins) and sacrificed at the times shown. Epididymal sperm were recovered and assessed for **a)** Total motility **b)** Progressive motility or **c)** Morphology. **d)** Shows a phase contrast image of the different types of sperm morphology seen 3 weeks post heat, compared to control (upper left). P value calculated by student’s t test, *=<0.05, N=3 shown for every time point.

### Testicular hyperthermia disrupts gene expression in round spermatids and spermatocytes

To investigate the transcriptional impact of testicular heat stress, mice were assigned to either a testicular hyperthermia (42°C for 30 minutes, N=5) or an anaesthetic control group (33°C/30 mins, N=5). After recovering from anaesthesia, mice were immediately sacrificed and round spermatids were isolated by Fluorescence-Activated Cell Sorting (FACS) as outlined by Kumar *et al,.* 2016 [49]. Although we aimed to capture the “immediate” transcriptional effects of heat stress, it is worth noting that the entire process – from the start of heating to obtaining an enriched cell population – took approximately 6 hours and attempts to reduce this timeframe proved unsuccessful. Purity assessments using H33342 stained nuclei showed ∼90% purity in round spermatids and (Supplementary Fig. 1). Using STA-PUT velocity sedimentation we were able to collect round spermatids at a purity comparable to FACS, as well as allowing isolation of spermatocytes at 80% purity in our hands.

Total RNA was extracted and both concentration and purity evaluated by a nanodrop spectrophotometer. Having passed quality control, RNA sequencing was performed and differential gene expression analysis was conducted DESeq2 (*p*<0.05 and ≥1.5 log2-fold change). PCA and feature mapping plots are provided in Supplementary Fig. 2. This analysis revealed 124 transcripts were upregulated while 10 were downregulated. This comparatively small number of gene changes is likely reflective of the idea that our experimental design aims to profile the earliest transcriptional response following *in vivo* testicular hyperthermia in isolated cells. The full list of differentially expressed transcripts is provided in Supplementary Table S1. Overall, this data shows a 93% bias towards increased transcript abundance, which is typically not seen in transcriptomic data. A Volcano plot showing the trend towards upregulation can be seen (Fig. 2a).

**Figure 2.**
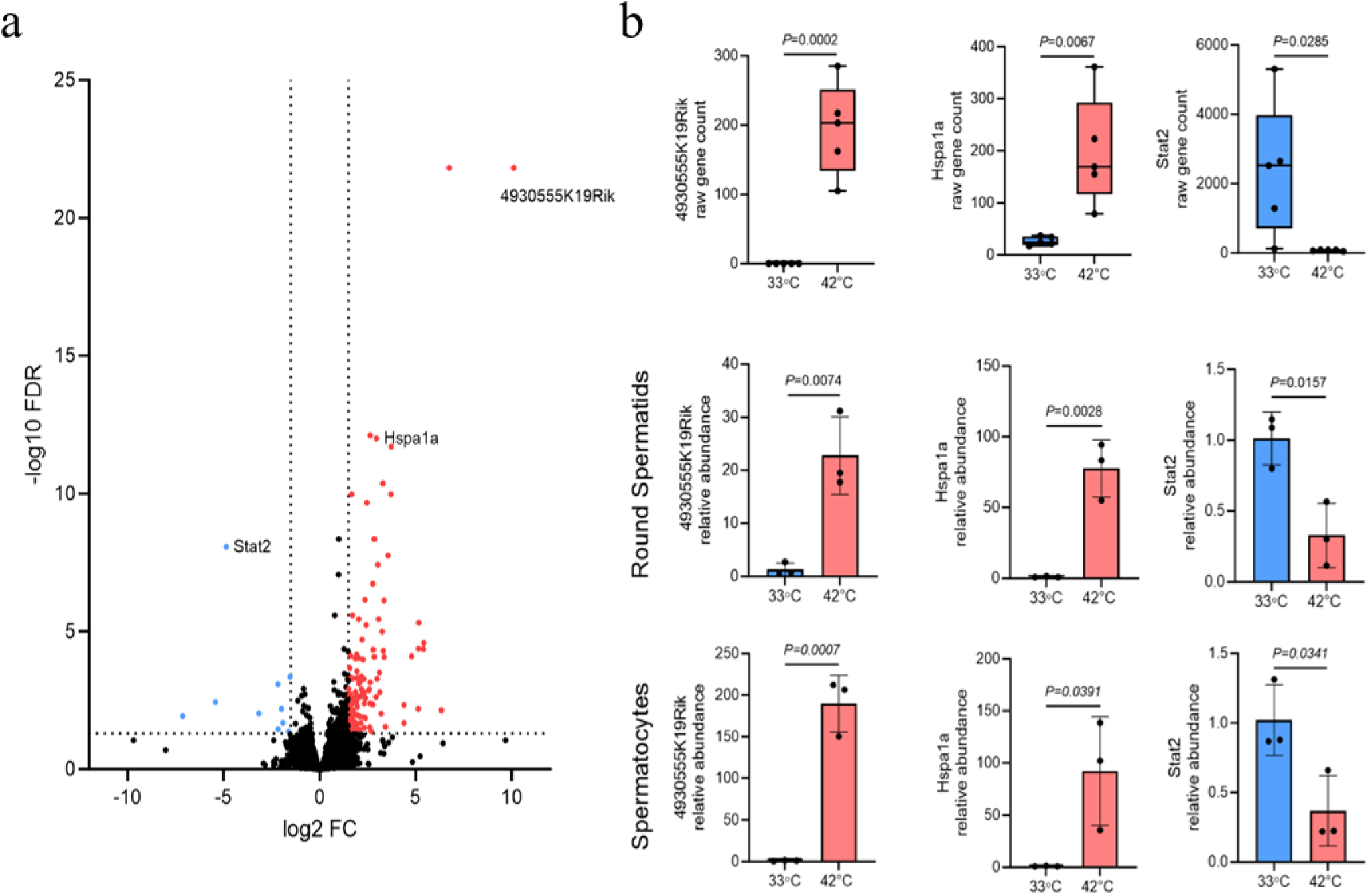
Differential gene expression in response to testicular heat stress. **a** Volcano plot of differentially expressed genes in heat stressed round spermatids (N=5, FDR >0.05, and absolute fold change >1.5). **b** Raw gene counts of genes selected for orthogonal confirmation. **c** RT-qPCR confirmation of differential gene expression in heat stressed round spermatids and **d** spermatocytes (N=3). P value calculated by student’s t test.

To validate our dataset, we performed RT-qPCR on isolated round spermatids by STA-PUT density gradient. Coincident with this, we collected enriched spermatocyte fraction, with the initial intent to see if gene changes were conserved amongst the male heat sensitive cell types. Three transcripts were selected based on their abundance and fold change. This included the most significant changing transcript LncRNA4930555K19Rik, which is enriched in the testis and of unknown function, *Hspa1a* a common heat shock response reporter gene, and *Stat2*, one of the few downregulated transcripts. Gene counts across 5 biological replicates from NGS data are shown for all three transcripts (Fig. 2b). Consistent with our sequencing data, we confirmed testicular hyperthermia led to an increase in LncRNA4930555K19Rik and *Hspa1a* expression, along with a decrease in *Stat2*, in round spermatids (Fig. 2c). Of interest, all three transcripts were shown to significantly change in enriched populations of heat stressed spermatocytes (Fig.2d). In the case of LncRNA4930555k19Rik and *Hspa1a*, a greater fold-change was observed in spermatocytes compared to round spermatids, suggesting that while spermatocytes and round spermatids are distinct cell types, they exhibit common transcriptional responses towards testicular hyperthermia.

### Little overlap in differentially expressed genes exists between our and other studies at the same timepoint post heat stress

To contextualise our findings, we compared our list of differentially expressed genes within heat stressed round spermatids with a recently published dataset that analysed transcriptional changes using whole testis RNA extracts following a comparable hyperthermia model (43°C, 30 minute water bath[32] ). Despite the shared stress conditions and analysis at similar timepoint (6 hours post heat stress), only one gene, *Hspa1a*, was upregulated across both datasets. This gene exhibited an 8.5 (log2)-fold increase in the whole testis dataset and a 3 (log2)-fold increase in our isolated round spermatid data. A full comparison of both datasets is provided in Supplementary table S2. The limited overlap in regulated genes likely results from differences in source material, which included whole testis RNA by others[32], versus RNA from isolated round spermatids by us.

### Testicular heat stress leads to changes in alternative splicing, albeit less consistent

Our sequencing protocol, with a read depth of 56-148 million reads per sample, enabled us to detect transcriptional changes beyond gene abundance, to include Local Splice Variations (LSVs). Using MAJIQ and VOILA, we identified 395, LSV events (Δψ>0.1, confidence interval >0.9) following heat stress in isolated round spermatids (Fig. 3a supplementary table S3). Interestingly, we observed notable inter-group variability in splicing patterns. For example, exon 5 of LncRNA4932415M13Rik was preferentially skipped in all heat stressed animals. However, in the control group, four out of five mice exhibited preferential exon inclusion, whereas one control mouse displayed a splicing pattern similar to those of the heat stressed group (Supplementary Fig. 3). This variability suggests inherent differences within individual mice exists. To validate LSV events, we conducted RT-qPCR using primer pairs specific to each condition of the variants within spermatids and spermatocytes. Three representative LSV events are shown as Sashimi plots of exon spanning reads (Fig. 3). In the case of *Fads2*, testicular hyperthermia led to an increase in intron retention in both round spermatids (Fig. 3b) and spermatocytes (Fig. 3c) when normalized to *β-actin* and an internal control targeting a conserved region of the *Fads2* transcript itself, consistent with the NGS data (Fig. 3a). Increased exon 5 skipping within LncRNA4932415M13Rik was identified in NGS data (Fig. 3d). Although RT-qPCR showed a comparable trend when normalised to *β-actin*, the change was not significant in either round spermatids (Fig. 3e), or spermatocytes (Fig. 3f). Similarly, in *Tas1r1* a decrease in intron retention was observed in heat stressed round spermatid NGS data (Fig. 3g) and validated by RT-qPCR when normalised to both *β-actin* and an internal control (Fig. 3h). This LSV event was absent in spermatocytes (Fig. 3i), suggesting a cell type specific response. Together this data suggests hyperthermia can induce LSV, and despite the fact that both *Fads2*[56]and *Tas1r1*[57] play an essential role in spermatogenesis GO analysis did not reveal functional pathway enrichment among LSV affected genes (padj>0.05). Additionally, the observed variability within the control group indicates that the identified LSVs alone are insufficient to explain the heightened sensitivity of germ cells to hyperthermia.

**Figure 3.**
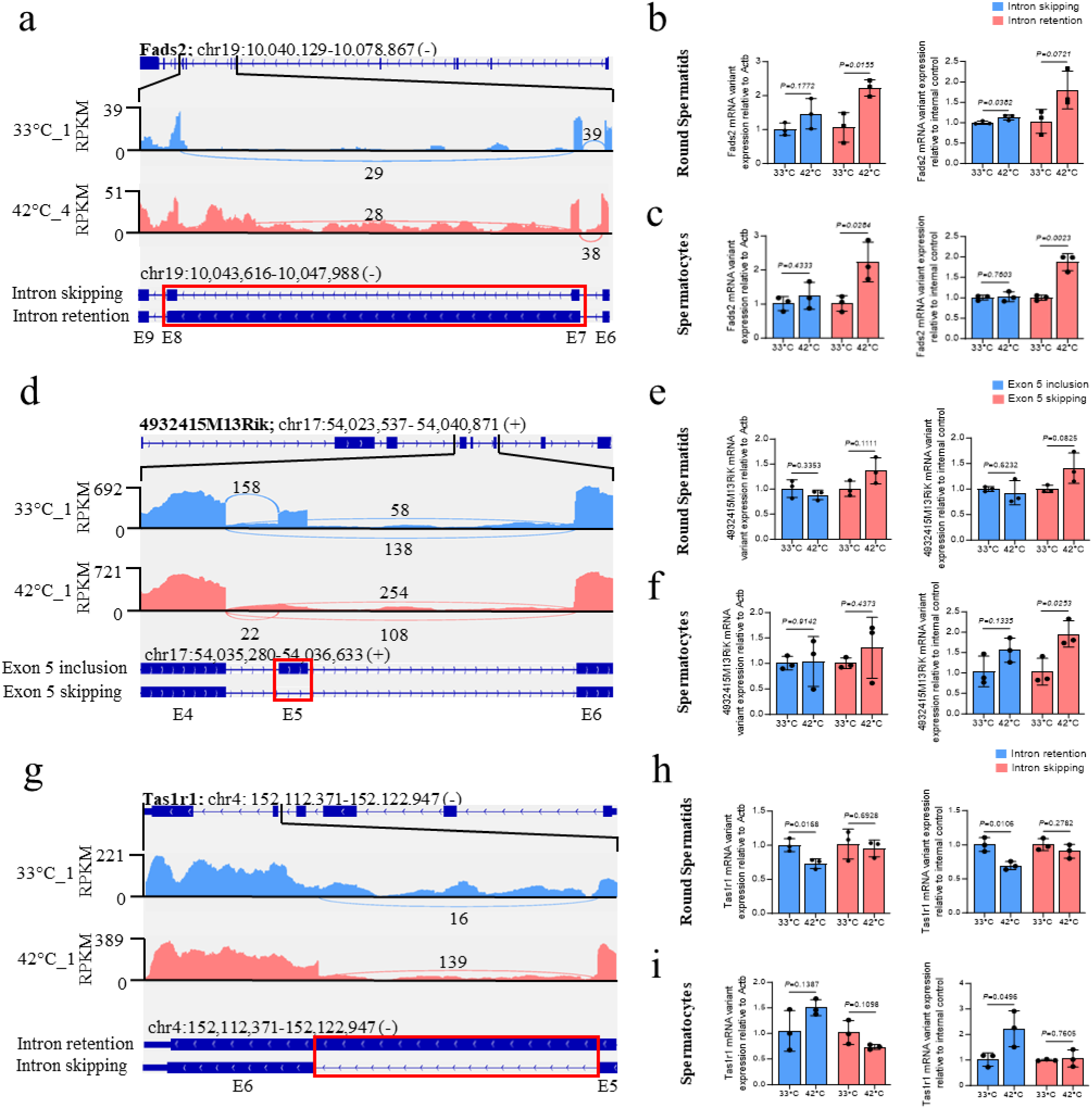
Local Splice Variation (LSV) within heat stressed spermatogenic cells. **a, d & g** Sashimi plots of representative LSV events identified within heat stressed round spermatids using MAGIQ (Δψ>0.1, confidence interval >0.9). RT-qPCR confirmation of LSV within heat stressed round spermatids (**b, e, & h)** and spermatocytes (**c, f, & i**, N=3). Transcript abundance relative to actin (left) and internal controls (right) are shown. P value calculated by student’s t test.

### Changes in alternative polyadenylation occurring following testicular heat stress

Alternative polyadenylation (APA) is known to occur in response to cellular stress and can impact cell fitness.[58–62] To investigate whether testicular hyperthermia affects polyadenylation, we analysed differential polyadenylation site usage between heat stressed and control round spermatids. We identified differential expression of 20 proximal polyadenylation usage (PPAU) sites (Fig. 4a) and 41 distal polyadenylation usage (DPAU) sites (Fig. 4b). However, many of these sites had low read counts or low percentage usage change between groups. Consequently, we opted not to validate APA changes using RT-qPCR, concluding much like LSV, APA are unlikely to be a driver of testicular heat stress. The full dataset of APA changes can be found in Supplementary table S4.

**Figure 4.**
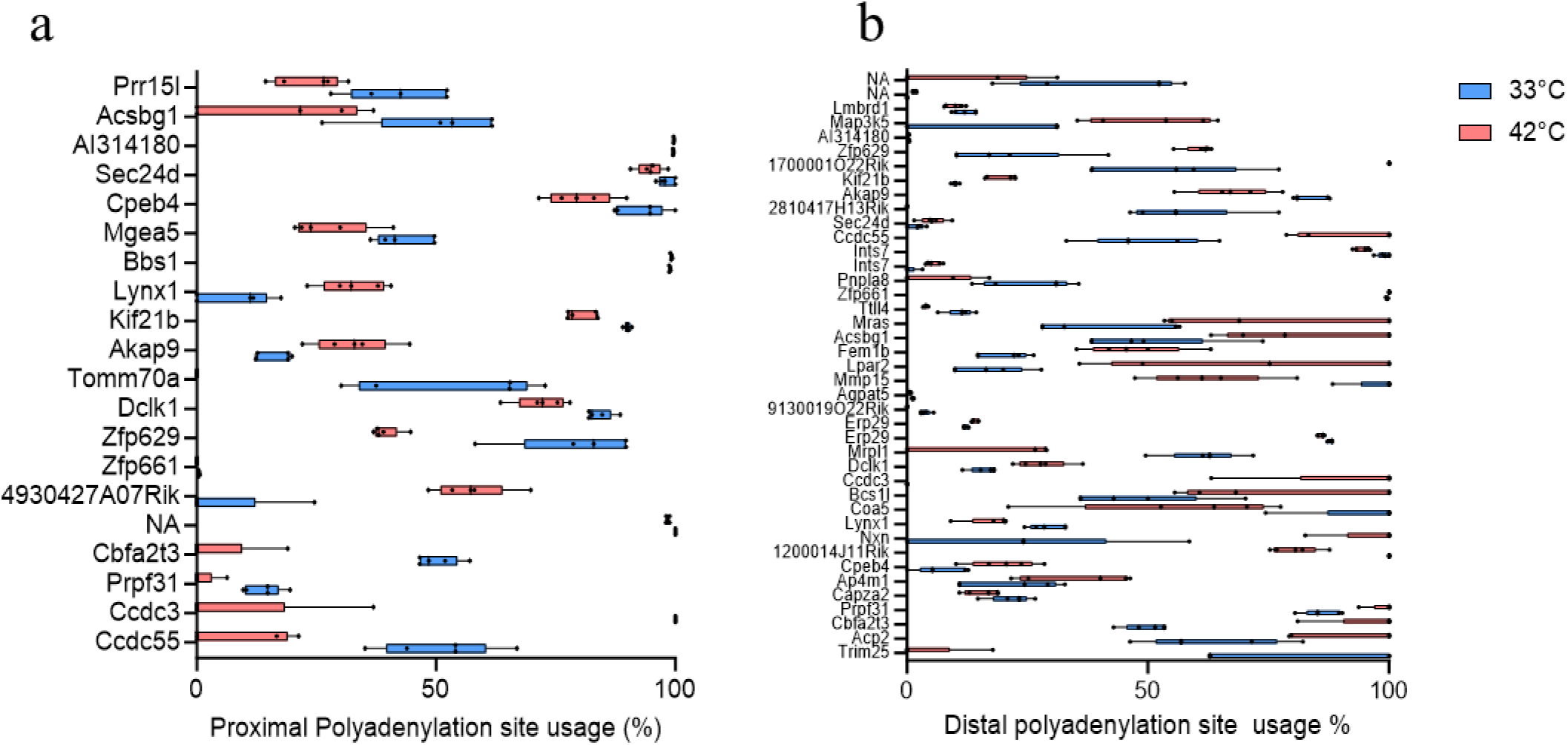
Alternative polyadenylation (APA) in heat stressed round spermatids. Differential percentage usage of **A** proximal polyadenylation sites and **B** distal polyadenylation sites within heat stressed round spermatids (N=5, padj>0.05). APA sites identified with QAPA.

### Testicular hyperthermia increases in Transposable Element expression including LINE1

Our initial analysis showed that 93% of transcripts were upregulated following testicular hyperthermia (Fig. 2a). This is quite unusual for “omic” studies, where it is typical to see near equal numbers of genes being up and down regulated. Further, the increase in transcript abundance were unlikely to be explained by LSV and APA changes described above. As such we next decided to investigate the role of testis enriched non-coding RNA as a potential cause. To gain insight into which non-coding RNA could be involved, we took advantage of the data we already obtained and reinvestigate it for the TE. Standard transcriptomic pipelines often overlook TEs during alignment of sequencing data to a reference genome, instead focusing primarily on typical mRNA coding species. However, we reasoned that if piRNA were affected by hyperthermia, this would be reflected in changes in TE abundance. As such we aligned our sequencing data to include TE annotations. Similar to the pattern observed in gene expression, TE transcripts showed remarkable bias towards upregulation upon heat stress, visualised by volcano plot (Fig. 5a). While Benjamani-Hochberg correction identified only one statistically significant upregulated TE in round spermatids, the overall trend pointed towards increased TE expression. Expression of individual TEs and their respective fold chance are provided in Supplementary table S1. Notably, both class 1 TEs (including LINEs, SINEs, and LTR retrotransposons) as well as class 2 DNA transposons showed increasing expression following hyperthermia.

**Figure 5.**
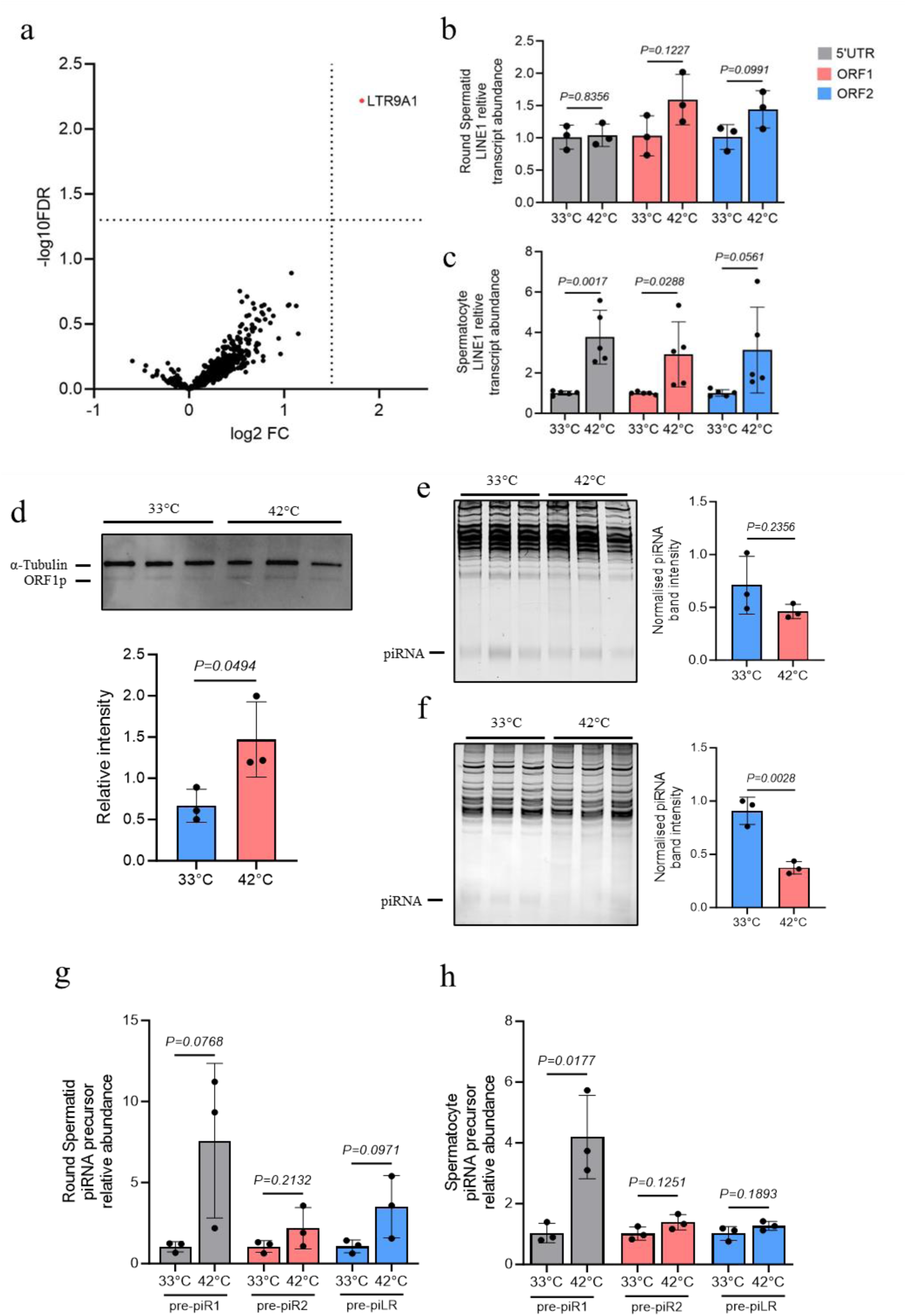
Increased Transposable element (TE) transcript abundance and decreased piRNA abundance 6H post testicular heat stress. **A** Volcano plot of differential TE expression within round spermatids 6 hours post testicular heat stress (N=5). **B** RT-qPCR found no significant increase in LINE1 transcript abundance within heat stressed round spermatids (N=3), but **C** significant upregulation in heat stressed spermatocytes (N=5). **D** Increasing ORF1p protein abundance in heat stressed spermatocytes as shown by western blot (Full image provided in Supplementary Fig S4). UREA-PAGE separation RNA followed by relative quantification of piRNA revealed no significant change within heat stressed round spermatids **E**, but a significant reduction in piRNA within heat stressed spermatocytes **F** (N=3). P value calculated by student’s t test. **G** qPCR was performed on 3 selected pre-cursor piRNA transcripts from either isolated round spermatids or **H** pachytene spermatocytes.

To validate these findings, we focused on LINE1, the most abundant and autonomously active TE in both the mouse and human genomes [63–65]. Using three primer sets targeting key regions of the LINE1 transcript (3’UTR, ORF1 and ORF2), we detected an average 3.3-fold increase in spermatocytes (Fig. 5c), while round spermatids exhibited an upwards (Fig. 5b), but not statistically significant trend in line with the NGS data. Active LINE1 encodes two proteins essential for template RNA binding and retrotranspostition, namely ORF1p and ORF2p respectively[63]. To determine the activity of upregulated LINE1 transcripts, we performed immunoblotting for ORF1p in heat stressed spermatocytes (Fig. 5d). This revealed a significant increase in ORF1p (∼45kDa) relative to α-tubulin, confirming increased translation of LINE1 after heat stress.

The bias towards upregulation of TE of all classes, alongside the observation that 93% of dysregulated gene transcripts were upregulated, suggested that piRNA (which normally supress transcripts) likely play a fundamental role. To investigate this further, we extracted total RNA from heat stressed and control precursor germ cells and quantified mature piRNA levels using 15% acrylamide denaturing Urea-PAGE and pre-cursor piRNA through qPCR. Spermatocytes exhibited a significant (41.14% ± 6.47% of control intensity) reduction in piRNA levels (Fig. 5f), which once again, correlates with the observed upregulation of LINE1mRNA and ORF1p. Interestingly, the loss of piRNA could not be explained by the loss of pre-cursor piRNA, which showed either no (Fig. 5g) or in one case a significant increase with spermatocytes following heat stress (Fig. 5h).

### Cells rebound 24-hours post heat stress

Previous transcriptional studies focus on later timepoints (24 hours post water bath, 3-5 days of cryptorchidism) and unlike our findings, they report relatively equal numbers of upregulated and downregulated piRNAs [33, 66]. To better characterize this response, we quantified piRNA levels and LINE1 abundance in enriched populations of spermatocytes and round spermatids at 24 hours post heat stress. Our data shows a partial recovery of global piRNA levels in spermatocytes, from 41.14% ± 6.47% of control intensity at 6 hours, to 56.76% ± 22.61% of control intensity at 24 hours (Fig. 6b). In round spermatids, while a reduction in piRNA band intensity was observed at 24 hours (Fig. 6a), it did not reach statistical significance at either timepoint. Interestingly, we noted a significant loss of LINE1 transcript abundance 24 hours after heat stress in round spermatids (Fig. 6c), and spermatocytes (Fig. 6d), a complete reversal of the upregulation seen at 6 hours (Fig. 5b). This unexpected decrease suggests a dynamic, time dependant regulation of TE in response to heat stress. Early during testicular hyperthermia there appears to be a loss of piRNA and consequent increase in TE levels. However, the cell appears to rapidly respond and compensate for this by upregulating piRNA, such that by 24 hours there is a loss in LINE1 transcript abundance.

**Figure 6.**
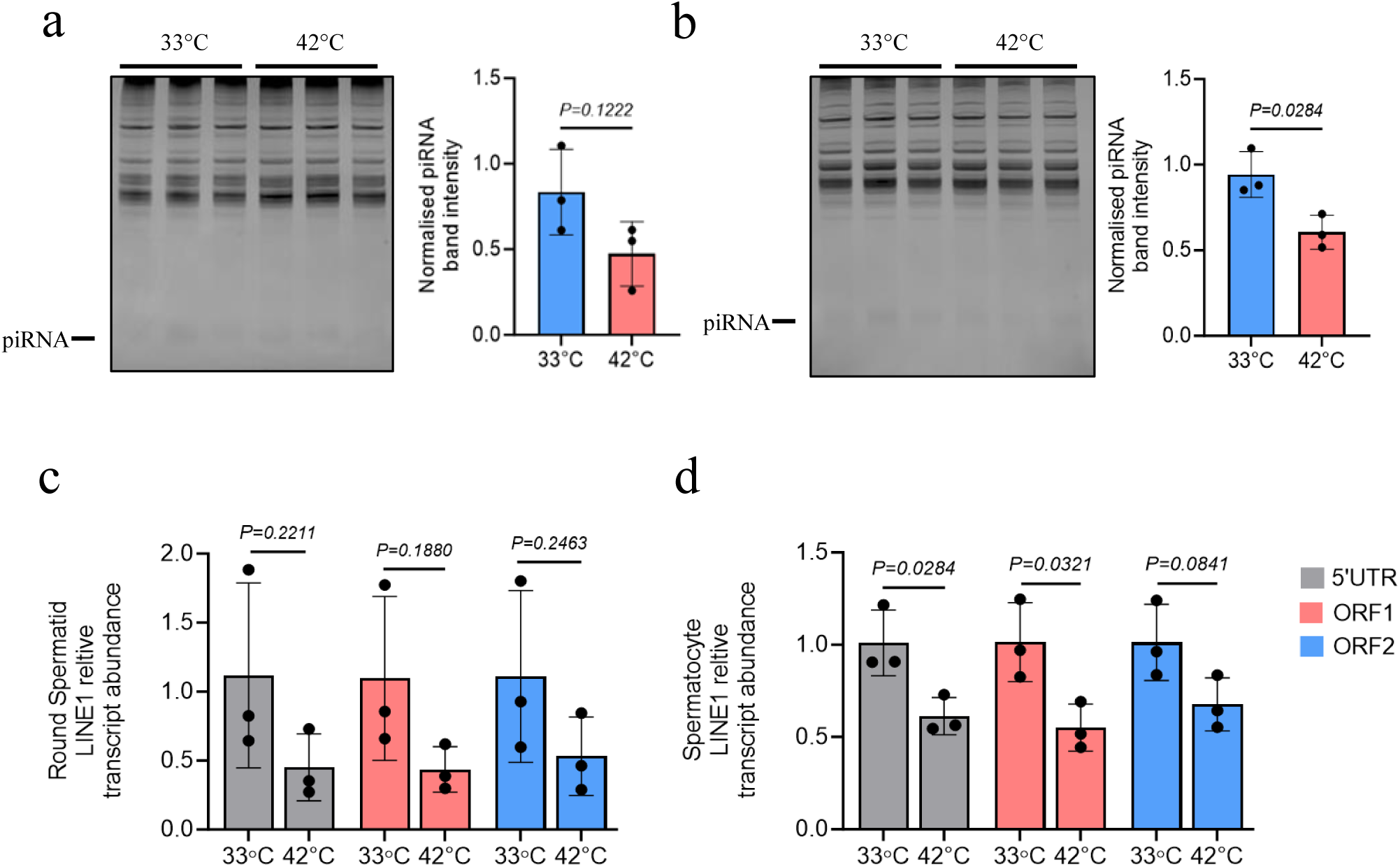
piRNA and LINE1 transcript abundance 24 hours post testicular heat stress. piRNA levels do not significantly differ from control in **A** round spermatids or **C** spermatocytes at 24 hours post testicular heat stress (N=3). LINE1 transcript abundance is reduced in both **C** round spermatids and **D** spermatocytes at 24 hours post heat stress (N=3). P value calculated by student’s t test.

### LINE1 expression is correlated with precursor germ cell DNA damage

The upregulation of LINE1, particularly in spermatocytes, prompted us to ask the question of whether this might underly the DNA damage seen in response to testicular heating. To assess this, we performed immunohistochemical analysis of testicular sections. The percentage of TUNEL positive cells as a proportion of total cells was significantly increased (0.12% ± 0.17 to 9.73% ± 3.86, N=3) 24 hours post heat stress (Fig. 7b), consistent with previous findings using similar water bath models[27, 30, 67–70]. Upon examination, it was evident that DNA damage was primarily among spermatocytes, based on nuclear morphology and location within tubules, consisted with the up-regulation of LINE-1.

**Figure 7.**
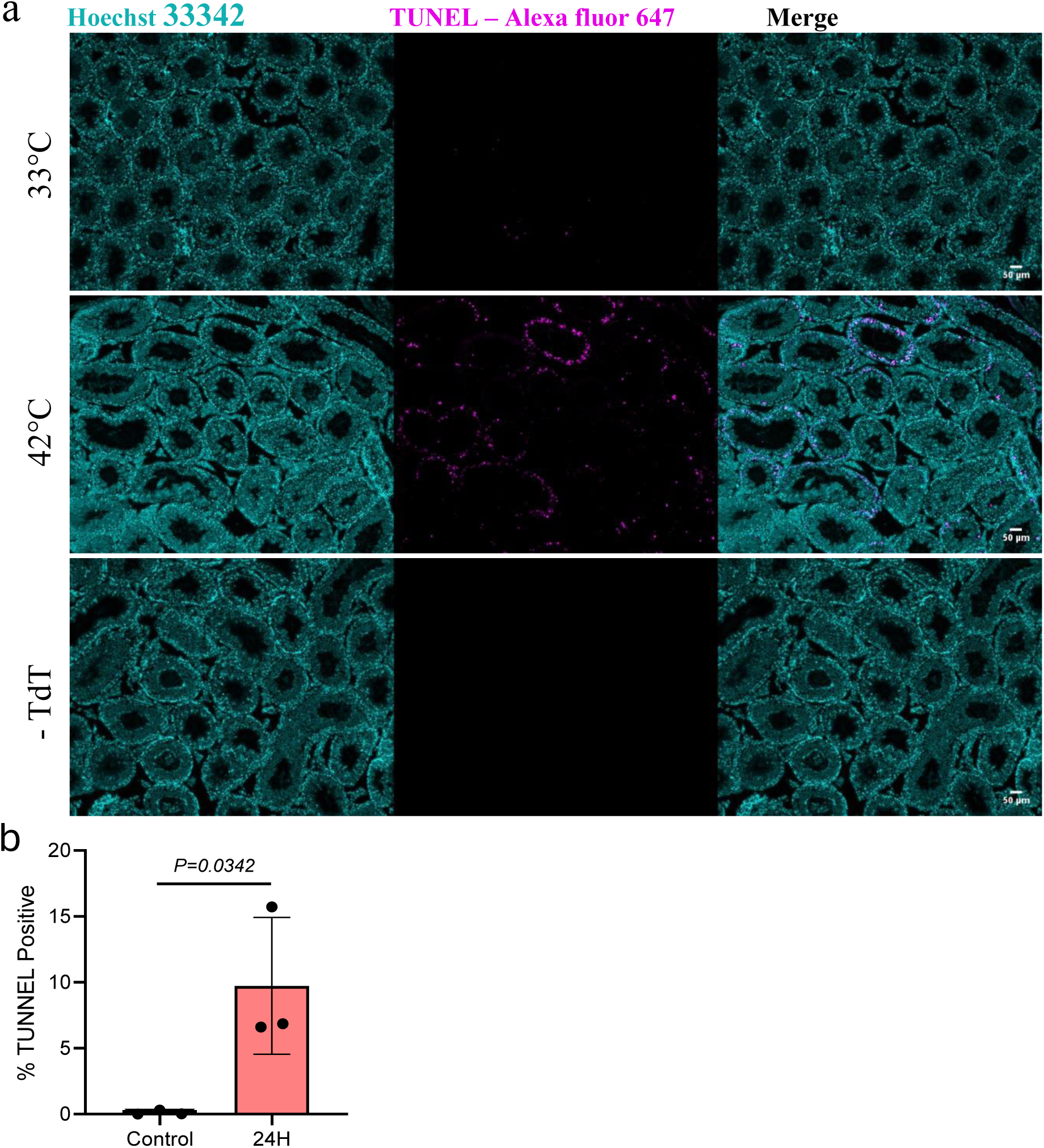
Increased spermatogenic cell DNA damage 24H post testicular heat stress. **A** TUNEL assay performed on testicular sections taken 24H post 30 min 33°C (control) and 42°C (heated) water bath (N=3). Nuclei counterstained with Hoechst 33342. **B** Percentage TUNEL positive cells increased 24 H post testicular heat stress. P value calculated by student’s t test.

## Discussion

Spermatogenesis is a remarkably unique process in that for optimal sperm production, the testis must remain lower than core body temperature. The detrimental effects of elevated testicular temperature have been recognised since 1893, when it was first linked to the loss of sperm production[71]. The molecular basis of testicular heat stress must account for three pertinent observations. Firstly, immunohistochemical analysis suggests spermatocytes are most sensitive to hyperthermia, followed by round spermatids[23]. Secondly, the mechanisms underlying heat sensitivity must be largely absent in somatic cells, which function optimally at core body temperature. Finally, any proposed mechanism must explain why testicular hyperthermia results in not only reduced sperm production but also elevated DNA damage and poor semen quality.

We propose that a rapid loss of piRNA fulfills all three criteria. Firstly, our data demonstrates that spermatocytes show a more pronounced reduction in piRNA levels and an increase in LINE1 abundance compared to round spermatids rapidly following hyperthermia. Secondly, piRNA are most abundant in the testis, wherein they are essential for normal sperm production, with the loss of piRNA biogenesis machinery consistently resulting in male infertility [37, 72]. Finally, the up-regulation of TE likely explain increased DNA damage.

Of note, a total loss of piRNA typically leads to azoospermia, however, the mouse model used here leads to impaired sperm quality 3 weeks post hyperthermia. We suggest this discrepancy is due to a critical threshold of piRNA. In our case, it is evident some piRNA remain at six hours post heat stress, with a trend to recovery after 24 hours. Residual piRNA may allow cells to progress through spermatogenesis, albeit producing lower quality sperm. This interpretation aligns with observations in cryptorchid models, where after 3-5 days of cryptorchidism piRNAs are dysregulated, but remain in sufficient number as to not completely inhibit spermatogenesis[33].

How then does hyperthermia lead to a loss of piRNA production? It is plausible that hyperthermia inhibits the translation biogenesis machinery, potentially through EIF2a phosphorylation and stress granule formation as reported by others[73]. However, in our model, following heat stress we see no change in the level of MIWI protein levels (data not shown) suggesting this in unlikely to be the case. An alternative hypothesis would suggest that hyperthermia affects the regulation of myeloblastosis oncogene-like 1 (MYBL1), the primary transcription factor responsible for piRNA production in mice[74]. Indeed, our greatest up- regulated transcript (*4930555K19RIK)* also has a MYBL1 protomer. Additionally, qPCR analysis of three pre-piRNA precursors revealed a significant increase in one case. These findings suggest two possible scenarios: (1) MYBL1 activity is dysregulated by temperature, leading to aberrant *4930555K19RIK* and pre-piRNA expression, or (2) the loss of mature piRNA results in the derepression of transcripts such as *4930555K19RIK*, which would normally be suppressed by piRNA. Further work is needed to clarify which interpretation is correct.

While we do not yet understand why there is a loss of piRNA following testicular hyperthermia, the downstream consequences are quite dramatic. Indeed, if sperm, that have developed under heat stress conditions, are used for fertilization, there are notably higher and significantly increased rates of embryo loss[12, 19–22]. Two key factors that likely contribute to this event. Following loss of piRNA, testicular hypothermia leads to heightened expression and activity of TE. While low TE activity during spermatogenesis is tolerated and can contribute to genetic diversity, unchecked activity poses risks of DNA damage and epigenetic changes[38–43, 75, 76].. As such, this increases the “chance” of genome instability and mutations occurring within protein coding genes. Secondly, piRNAs not only prevent translation of transcripts through MIWI (PIWIL1), but also permanently inhibit transcription of many genes (notably TEs) by directing the methylation of promoter sites[75]. In this scenario, DNA-methylation appears to be continual process occurring throughout spermatogenesis whereby genes are switched either on or off, including the spermatocyte and spermatid stage of development [77]. Typically, MILI (PIWIL2) binds piRNAs that are derived from mRNA products including TEs. The MILI- piRNA complex then translocates to the nucleus. Herein histone methyltransferase DNMT3C is recruited to target promoter regions via piRNA homology recognition[78]. This leads to hypermethylation of the gene thereby inhibiting transcription[78]. If hyperthermia leads to a loss of piRNA, the methylation of genomic sites may not be established correctly during spermatogenesis which has the potential for epigenetic consequences and higher rates of embryo loss among offspring.

Overall, our work shows that testicular hyperthermia is not just detrimental for sperm production, but also for genome integrity. Testicular heat stress leads to a rapid loss of piRNA and upregulation of TE. In the case of spermatocytes there is also an increase in DNA damage. This has implications not only for male fertility, but also likely leads to genomic abnormalities and epigenetic changes with consequences for offspring if such cells are used for fertilisation.

