## Supplementary Figures S1-S4 for "Acute testicular hyperthermia leads to a rapid loss of global piRNA levels and a consequent increase in transcript abundance, including LINE1 activity within heat-sensitive male germ cells"

**Supplementary Fig. 1.** Representative images of **A** whole testis single cell suspension, **B** isolated round spermatid (arrow) and **C** isolated spermatocyte (star) populations following STA-PUT velocity sedimentation. Nuclei stained with Hoechst 33342 for determination of cell type based on nuclear morphology.

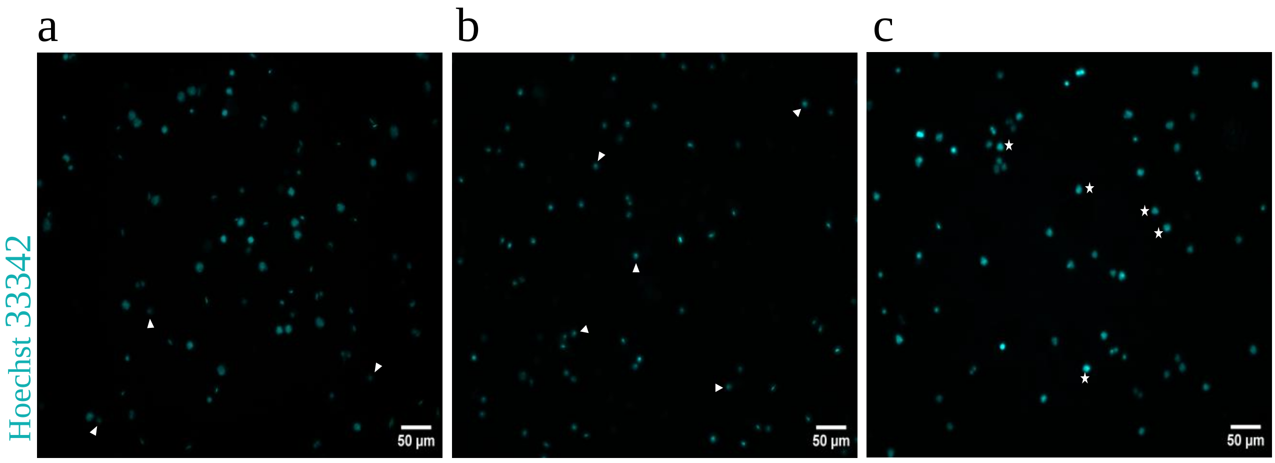

**Supplementary Fig. 2. A** Principal component analysis of top 500 variable genes shows separation of experimental groups. **B-D** Percentage counts mapped to TE , mitochondrial genes and genes. **E** Scatterplot of number of mapped reads, number of features detected, and percentage of features mapped to mitochondrial genes per sample.

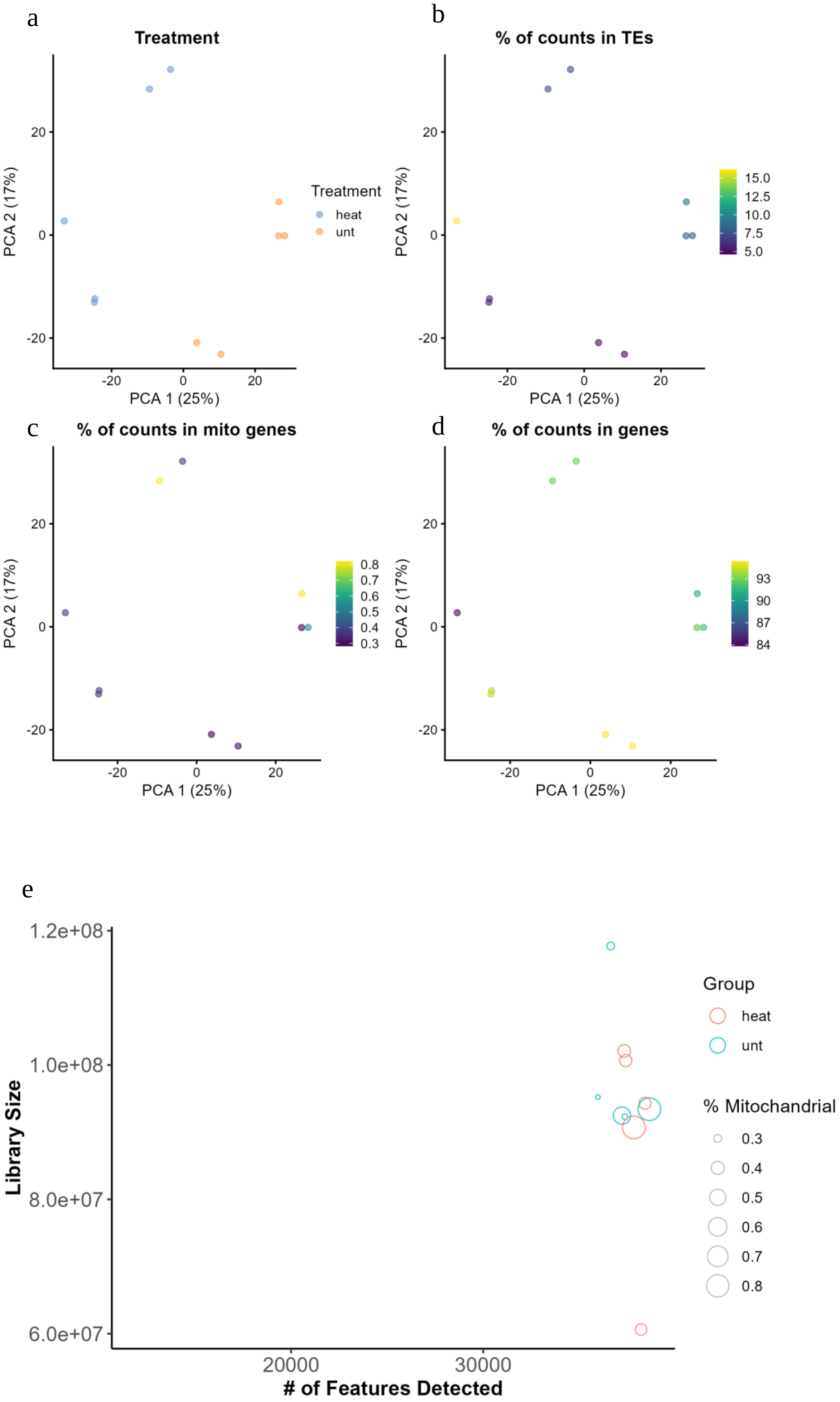

**Supplementary Fig. 3.** Sashimi plot of LSV events selected for orthogonal confirmation by qPCR. Selected events **A** gene-Fads2:s:10047636-10047712 **,B** gene-4932415M13Rik:s:54035280-54035523 with visible control group intervariability, and **C** gene-Tas1r1:s:152115201-152115361. Plots generated by Interactive Genomics Viewer (IGV).

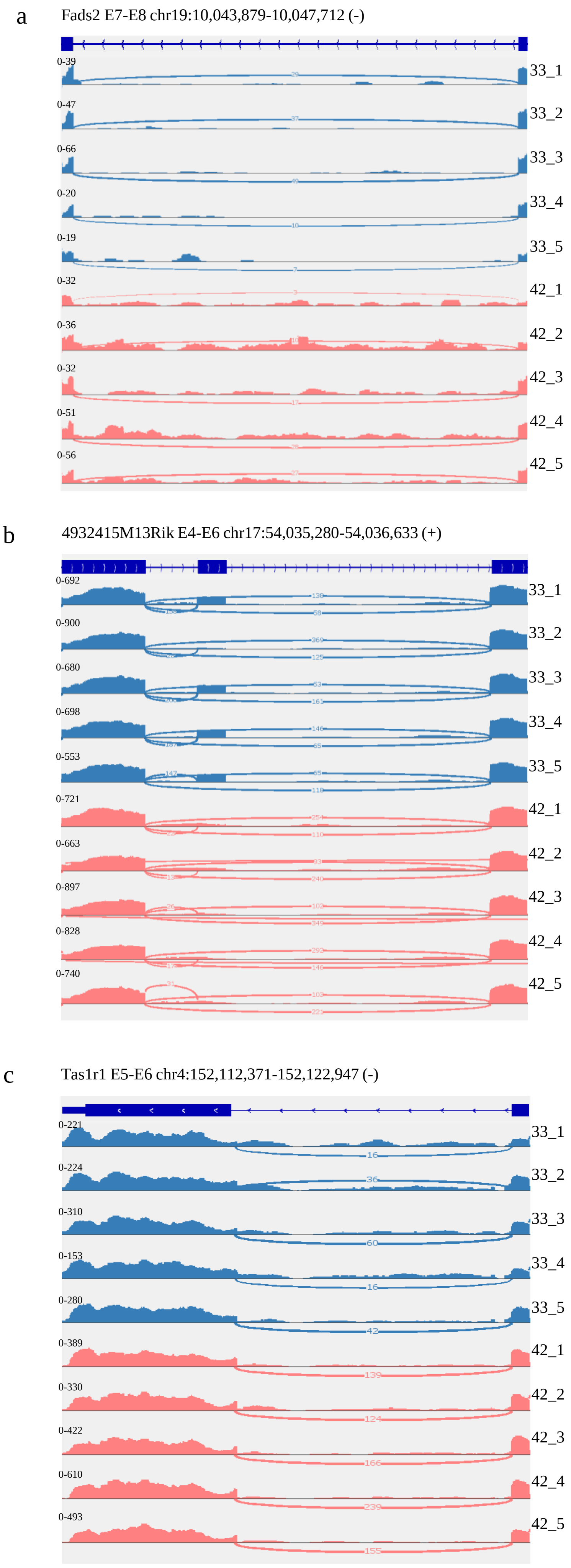

**Supplementary Fig. 4.** Relative ORF1p protein abundance increases within heat stressed spermatocytes (N=3), relevant MW markers shown.

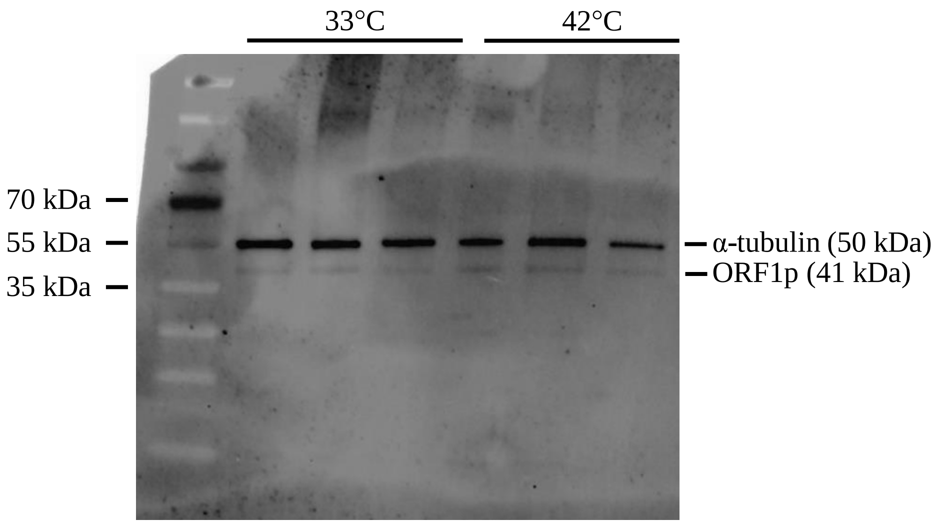
